# Convergence of cortical types and functional motifs in the mesiotemporal lobe

**DOI:** 10.1101/2020.06.12.148643

**Authors:** Casey Paquola, Oualid Benkarim, Jordan DeKraker, Sara Lariviere, Stefan Frässle, Jessica Royer, Shahin Tavakol, Sofie Valk, Andrea Bernasconi, Neda Bernasconi, Ali Khan, Alan Evans, Adeel Razi, Jonathan Smallwood, Boris Bernhardt

## Abstract

The parahippocampus-hippocampus complex in the mesiotemporal lobe (MTL) is implicated in many different cognitive processes, is compromised in numerous disorders, and exhibits a unique cytoarchitectural transition from six-layered isocortex to three-layered allocortex. Our study leveraged an ultra-high-resolution histological reconstruction of a human brain to (i) develop a continuous surface model of the MTL iso-to-allocortex transition and (ii) quantitatively characterise the region’s cytoarchitecture. We projected the model into the native space of *in vivo* functional magnetic resonance imaging of healthy adults to (iii) construct a generative model of its intrinsic circuitry and (iv) determine its relationship with distributed functional dynamics of macroscale isocortical fluctuations. We provide evidence that the most prominent axis of cytoarchitectural differentiation of the MTL follows infolding from iso-to-allocortex and is defined by depth-specific variations in neuron density. Intrinsic effective connectivity exhibited a more complex relationship to MTL geometry, varying across both iso-to-allocortical and anterior-posterior axes. Variation along the long axis of the MTL was associated with differentiation between transmodal and unimodal systems, with anterior regions linked to transmodal cortex. In contrast, the iso-to-allocortical gradient was associated with the multiple demand system, with isocortex linked to regions activated when task demands prohibit the use of prior knowledge. Our findings establish a novel model of the MTL, in which its broad influence on neural function emerges through the combination micro- and macro-scale structural features.

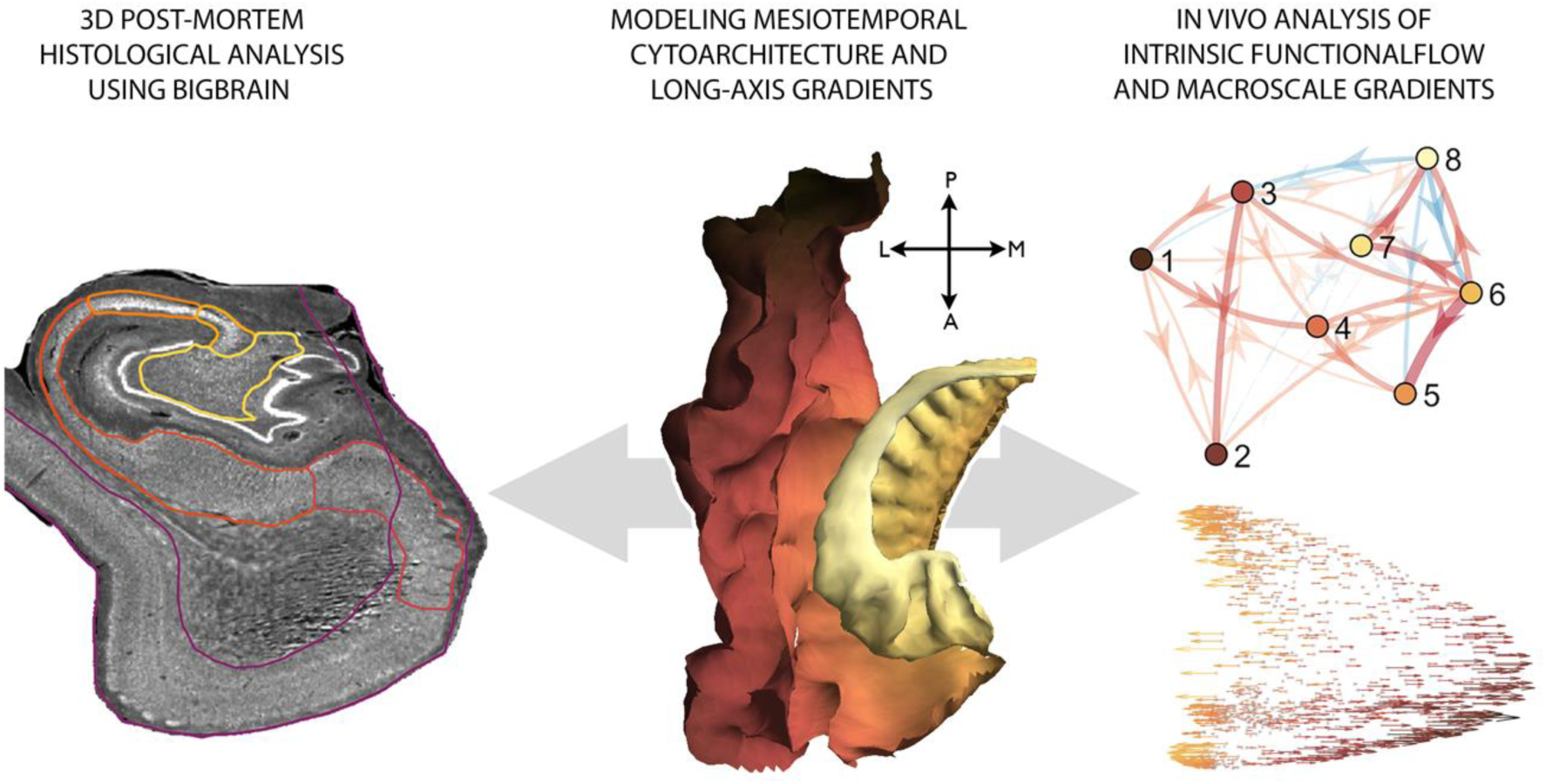

## Introduction

The mesiotemporal lobe (MTL) is implicated in a diverse range of cognitive processes, notably memory, navigation, socio-affective processing, and higher-order perception ^1,2,11,3–10^. Contemporary views recognise that the broad role this region plays in human cognition emerges through interactions between neural codes within the MTL and the rest of the brain ^10,12–14^. Motivated by its role in spatial processing ^15^, it has been suggested that the MTL plays a key role in the formation of cognitive maps ^16^ by providing a low dimensional representational scheme of different situational contexts ^15,17^. A widely held assumption posits that the broad role of the MTL in human cognition results from both its intrinsic circuitry and position within the larger cortical architecture, which places the MTL at the apex of multiple hierarchies ^18–20^.

Unique in the mammalian brain, the mesiotemporal cortex folds in upon itself, producing the curled hippocampal formation that sits atop the parahippocampal region. This infolding is accompanied by a shift from six-layered isocortex to three-layered allocortex ^21^. During ontogeny, isocortex (from iso-“equal”) differs from allocortical (from allo-“other”) hippocampus, which lacks clear lamination prenatally ^22,23^. Such variations in cytoarchitecture are thought to be involved in the transformation from unimodal to transmodal information ^24^, as well as degree of synaptic plasticity low in iso-, and high in allocortex ^25^. The transition from iso-to-allocortex occurs in smooth streams of gradation emanating from the hippocampus ^22,26,27^. This wave of laminar differentiation is phylogenetically conserved ^22,28–30^ and classic histological studies suggested that it is accompanied by step-wise changes in neuronal density at various cortical depths ^31^. Nevertheless, compared to other mammals, the human MTL has exaggerated folding, more extensive lamination of the entorhinal cortex as well as a more prominent appearance of the Cornu Ammonis (CA) 2 subfield, highlighting the need to characterise the region’s microstructure in our species ^21,26,32^.

Substantial cytoarchitectural differentiation within the MTL holds important implications for signal flow. Degree of laminar differentiation is generally thought to reflect the origin and termination patterns of axonal projections ^33^, and so the connectivity between cytoarchitecturally distinct MTL subregions may thus be described as feedforward or feedback based on relative iso-to-allocortical position. Accordingly, feedforward projections flow from iso-to-allocortex, whereas feedback projections flow from allo-to-isocortex ^33,34^. Thus, mapping directional signal flow along the axis could inform upon the role of the MTL in feedforward or feedback processing. Furthermore, cytoarchitectural similarity is tightly coupled with likelihood of interareal connectivity, a principle known as the “structural model” ^33,35–37^, suggesting that the intrinsic circuitry of the MTL is guided by the iso-to-allocortical axis. Consistent with this view, key fibre pathways in the MTL, notably the perforant pathway, mossy fibres, and Schaffer collaterals, are typically defined by their relation to hippocampal subfields (subiculum, CA1-4, dentate gyrus) that are sequentially organised along the iso-to-allocortical axis ^38^. In addition to the iso-to-allocortical shifts that follow hippocampal infolding, neurobiological and functional properties of the MTL system also appear to be organised with respect to a second, anterior-posterior, axis ^38–42^. Tract-tracing studies in rodents have shown that anterior-posterior gradients determine hippocampal connectivity to entorhinal cortices ^40^ and functional neuroimaging studies in humans have illustrated distinct combinations of anterior-posterior and lateral-medial topographies in the entorhinal and parahippocampal cortex ^43,44^. Recently, data-driven analyses of resting state functional magnetic resonance imaging (MRI) have confirmed marked differentiation of anterior and posterior aspects of the hippocampus in terms of intrinsic functional connectivity ^45–47^.

Given the established role of the MTL as a hub of multiple large-scale networks ^48–51^, the combination of both iso-to-allocortical and anterior-posterior axes may underlie its’ broad contribution to neural function. One potential advantage of this architecture could be that operations implemented by the internal hippocampal circuitry, such as pattern separation and completion ^52–55^, could be realised across multiple cortical processing zones in a distributed and coordinated manner. In this way, the coarse representations in the MTL could be mirrored by neural motifs that are present across the broader cortical system. Contemporary accounts have shown that whole-brain functional organisation can be understood as discrete communities such as the sensory, attention, and transmodal networks ^56^ and that the relationships between these communities can be compactly represented by gradients of cortex-wide connectivity ^57^. Critically, these approaches highlight overlapping neural motifs that are thought to play an important role in cognitive function, with one functional gradient differentiating sensory from transmodal systems that harbor cognitive processes increasingly reliant on information from memory ^58,59^. A second large-scale functional gradient reflects the dominance of networks involved in complex tasks in which there is no predetermined schema with which to guide behaviour, also referred to as the multiple demand system ^60,61^. Our study explores the mapping between MTL architecture and associations to different functional gradients, to establish whether this relationship provides a plausible mechanism through which the MTL elicits a broad role in neural function.

We leveraged an ultra-high resolution 3D reconstruction of a histologically stained and sliced human brain [*BigBrain* ^62^] to build a continuous model of the MTL architecture. Surface based approaches were used to sample intracortical microstructure across multiple cortical depths along both the iso-to-allocortical and anterior-posterior axes. We hypothesised that the iso-to-allocortical axis would be the principle gradient of cytoarchitectural variation in the MTL and identified salient cytoarchitectural signatures of this transition using supervised machine learning. We then translated the histology-based model of the MTL to *in vivo* functional neuroimaging to assess how the iso-to-allocortical and anterior-posterior axes covary with intrinsic and large-scale network connectivity. Building upon established models of internal hippocampal circuitry ^38^, we hypothesised that the strongest signal flow throughout the MTL would follow the iso-to-allocortical axis, and tested directional signal flow via dynamic causal modelling of resting state functional MRI ^63^. In addition to testing consistency of these intrinsic circuit characteristics across the anterior-posterior axis, we assessed how both the iso-to-allocortical and anterior-posterior axes relate to macroscale functional organisation by assessing the topographic representation of large-scale functional gradients in the MTL space.

## Results

We developed a detailed model of the confluence of cortical types in the parahippocampus-hippocampus complex based on an 40μm 3D histological reconstruction of a human brain (see *Methods;* **Figure 1A**), which we then translated to *in vivo* functional imaging and macroscale connectomics. Considering MTL cytoarchitecture, we observed a strong gradient along the iso-to-allocortical axis, with depth-wise intracortical profiles acting as robust predictors of spatial axis location. The most predictive feature was consistent across the anterior-to-posterior regions, suggesting a preserved iso-to-allocortical architecture along the MTL long axis. Mapping the cortical confluence model to *in vivo* resting state functional MRI, intrinsic signal flow was strong along the iso-to-allocortical axis but interacts with the long-axis. We found that the iso-to-allocortical and anterior-to-posterior axes differentially contributed to distinct dimensions of macroscale functional systems, with anterior-to-posterior position reflecting trade-offs between sensory and transmodal function, and iso-to-allocortical positions related to the relative contribution of the multiple demand system.

**Figure 1.**
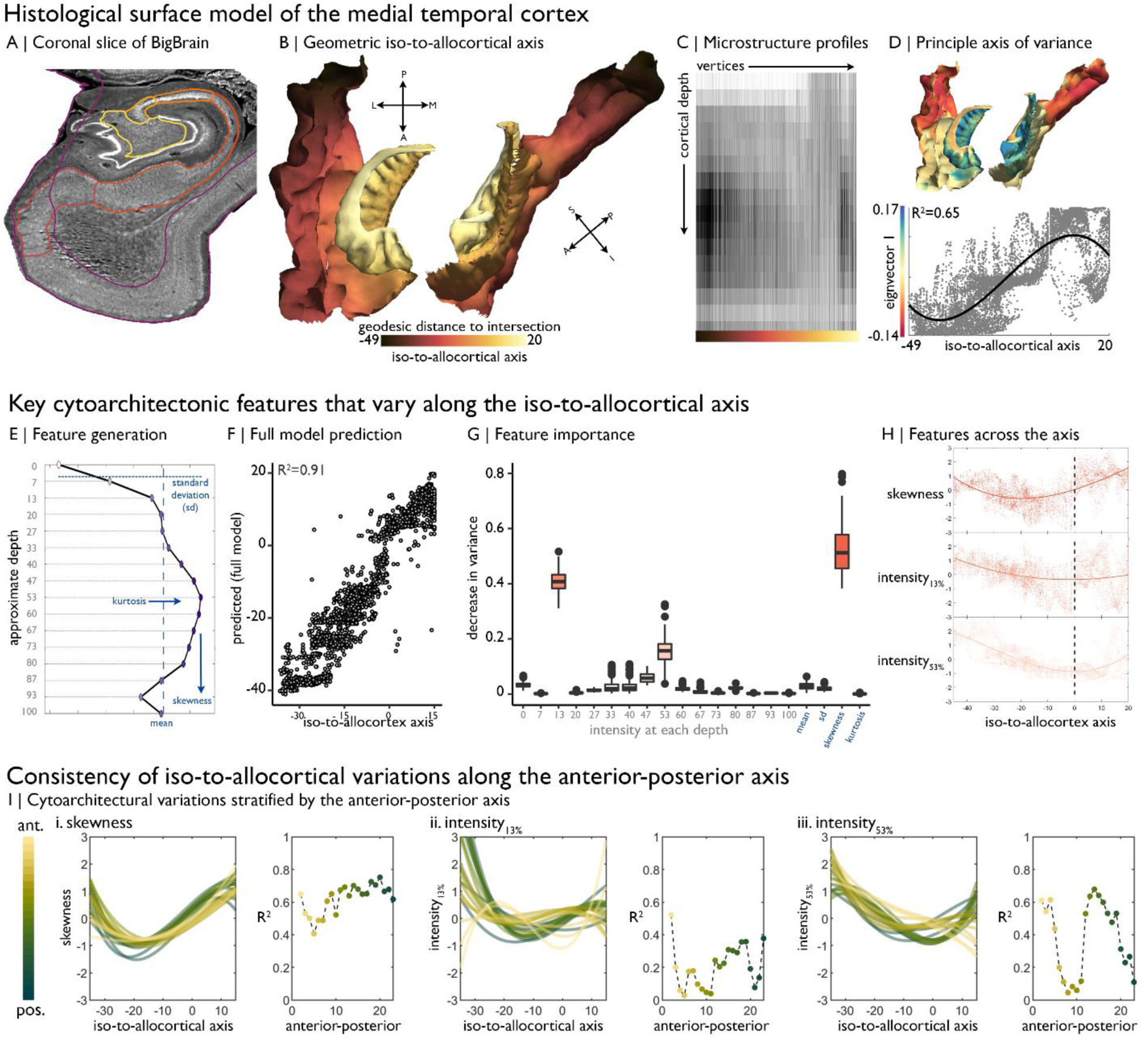
Cytoarchitecture profiling of the mesiotemporal confluence. **A)** Isocortical and allocortical surfaces projected onto the 40um BigBrain volume. Notably, conventional isocortical surface construction skips over the allocortex. Purple=isocortex. Red=subiculum. Dark orange=CA1. Orange=CA2. Light orange=CA3. Yellow=CA4. **B)** The iso-to-allocortical axis was estimated as the minimum geodesic distance to the intersection of the isocortical and hippocampal surface models. **C)** Intensity sampling along 16 surfaces from the inner/pial to the outer/white surfaces produced microstructure profiles. Darker tones represent higher cellular density/soma size. **D)** *Above*. The principle eigenvector/axis of cytoarchitectural differentiation, projected onto the confluent surface. *Below*. The association between the principle eigenvector and the iso-to-allocortical axis, with the optimal polynomial line of best fit. **E)** Cytoarchitectural features generated for each microstructure profile. **F)** Empirical *vs* predicted position on the iso-to-allocortical, based on supervised random forest regression with cross-validation. **G)** Feature importance was approximated as how much the feature decreased variance in the random forest. **H)** Cubic fit of each selected feature with the iso-to-allocortical axis. Feature values are z-standardised. **I)** Line plots show the cubic fit of cytoarchitectural features to the iso-to-allocortical axis within 23 bins of the anterior-posterior axis (*yellow-to-green*). Neighbouring scatter plots depict the goodness of fit (adjusted R^2^) of each polynomial, showing high consistency of the pattern of skewness.

### Cytoarchitectural analysis of the iso-to-allocortical confluence

We first mapped the iso-to-allocortical axis as the minimum geodesic distance of a vertex to the intersection of the isocortical and allocortical surface models (**Figure 1B**). To examine the cytoarchitecture of the cortical confluence, we generated microstructure profiles, reflecting staining intensity across 16 cortical depths (**Figure 1C**). Unsupervised manifold learning of microstructure profile data revealed that the principle gradient of cytoarchitectural differentiation, accounting for 15.6% of variance, closely approximates the geometric iso-to-allocortical axis (r=0.76, p<0.001; **Figure 1D**). This relationship appeared nonlinear (adjusted R^2^ for polynomials 1-3=0.58/0.59/0.65), and CA2 expressed a different cytoarchitecture than expected by its position on the geometric map (**Figure S1**). Lower eigenvectors were not spatially correlated with the iso-to-allocortical axis (0.01<|r|<0.31), demonstrating specificity of the first eigenvector in mapping this cytoarchitectural gradient (see **Figure S1**, for lower eigenvectors).

Microstructure profile features provided a highly accurate mapping of the iso-to-allocortical axis based on random forest regression (R^2^=0.91±0.005; **Figure 1E-F**). Profile skewness, a feature sensitive to depth dependent changes in staining intensity, as well as intensity at upper (∼13% depth) and mid surfaces (∼53% depth) were key features in learning the axis (average reduction in variance: 53/41/16%; **Figure 1G**). These three features alone accounted for most variance in out-of-sample data (R^2^=0.86±0.006; **Figure S2**). Microstructure profile skewness increased from iso-to-allocortex, which pertained to a shift from relatively low intensities at upper surfaces to a flatter microstructure profile (**Figure 1H**, see **Figure S2** for exemplar microstructure profiles and feature values). Intensity at ∼13% depth approximately corresponds to the layer1/2 boundary in the isocortex and exhibited a rapid uptick at the iso-allocortical intersection. Intensity at ∼53% depth approximately aligns with the peak intensity of the average microstructure profile, signifying high cellular density, and reached a minimum around the intersection of the iso- and allo-cortex.

Next, we sought to test whether the iso-to-allocortical variations were consistently expressed along the anterior-posterior axis. Independently inspecting 1mm coronal slices of the *post mortem* data, we found that microstructure profile skewness consistently increased along iso-to-allocortical axis, regardless of anterior-posterior axis (mean±SD adjusted R^2^=0.64±0.17, **Figure 1I *left***). On the other hand, depth-dependent intensities were more variable along the anterior-posterior axis (13% depth: goodness of fit=0.20±0.12; 53% depth: goodness of fit=0.40±0.24) and exhibited different iso-to-allocortical patterns in the anterior portion. Thus, while certain depth-dependent intensities systematically varied across both geometric axes, the most salient cytoarchitectural feature (skewness) consistently and gradually decreased across the iso-to-allocortical axis regardless of long axis position.

### Iso-to-allocortical gradients in internal signaling

Functional signal flow within axes of the mesiotemporal confluence was examined via dynamic modelling of resting state functional MRI, using a generative model of effective connectivity, after transforming the cortical confluence model first from histological to stereotaxic MNI152 space (**Figure 2A**), and then to native functional MRI spaces in a group of 40 healthy adults (see *Method*s for demographics, acquisition and preprocessing). In brief, we extracted blood oxygen level dependent timeseries from each voxel of the cortical confluence during a resting state scan, averaged timeseries within a discrete set of bins of the iso-to-allocortical axis and carried out Bayesian model reduction.

**Figure 2.**
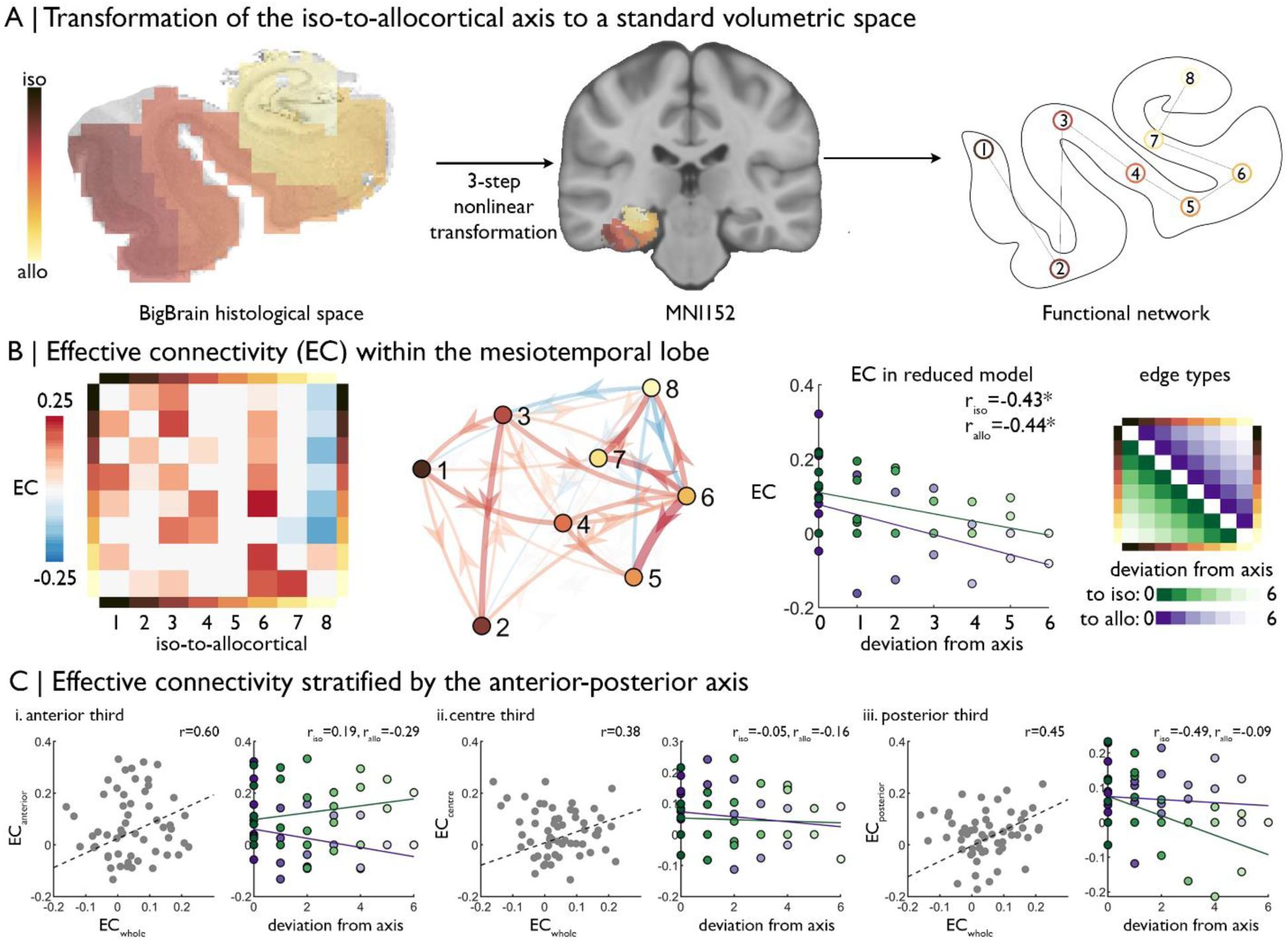
Intrinsic functional signal flow across the cortical confluence. **(A)** The surface-based iso-to-allocortical axis was projected into the native volume space of BigBrain, then registered to stereotaxic MNI152 space. We used this as a standard atlas to extract resting state functional MRI timeseries. An 8-node decomposition was used for the dynamic model (subregion overlap shown in Supplementary Figure 3). **(B)** *Left* Posterior estimates of effective connectivity (EC) between 8-nodes of the iso-to-allocortical axis. Columns are seeds and rows are targets. *Right*. Strength of effective connectivity with deviation from the iso-to-allocortical axis, stratified by direction. * signifies p_FDR_<0.05. **C)** Association of effective connectivity within anterior-posterior thirds of cortical confluence model (y-axes) with effective connectivity of the whole model and deviation from the iso-to-allocortical axis (x-axes).

This model showed dense, bidirectional effective connectivity across the MTL (**Figure 2B**). To test whether effective connectivity predominantly followed the iso-to-allocortical axis, and if so in which direction, we labelled edges by deviation from the axis, whereby lower deviation represented an edge more concordant with the iso-to-allocortical axis (**Figure 2B *far right*)**. Deviation from the iso-to-allocortical axis was related to lower effective connectivity in both directions (to isocortex r_iso_=-0.43, p=0.02; to allocortex: r_allo_=-0.44, p=0.02; **Figure 2B *right***). Furthermore, the final bin, containing CA4, inhibited other regions and the sixth bin, corresponding to the subiculum, was the strongest driver of excitatory effective connectivity (see **Figure S3** for subfield overlap).

Next, we tested the interaction of iso-to-allocortical and anterior-posterior axes in determining intrinsic signal flow by constructing the dynamic model within thirds of the anterior-posterior axis. If only the iso-to-allocortical axis determines the intrinsic connectivity, then the pattern of effective should be conserved within each third. Effective connectivity estimates, calculated from a model within thirds of the anterior-posterior axis were moderately correlated with effective connectivity estimates from the full mesiotemporal confluence model (r=0.60/0.38/0.45, **Figure 2C**). Unlike the full mesiotemporal confluence, however, we did not consistently observe inverse correlation between deviation from the iso-to-allocortical axis and effective connectivity in these spatially restricted models (**Figure 2C**).

### Iso-to-allocortical and anterior-posterior axes represent macroscale functional organisation

After demonstrating subtle interactions of iso-to-allocortical and anterior-posterior axes in cytoarchitectural and intrinsic signalling of the MTL, we examined the interplay of both axes in the topographic representation of macroscale functional systems within the MTL. To this end, we examined conventional rs-fMRI connectivity between MTL voxels and isocortical parcels, with large-scale networks being characterized with respect to both previously established canonical functional communities ^56^ and macroscale connectome gradients ^57^ (**Figure 3A-C**).

**Figure 3.**
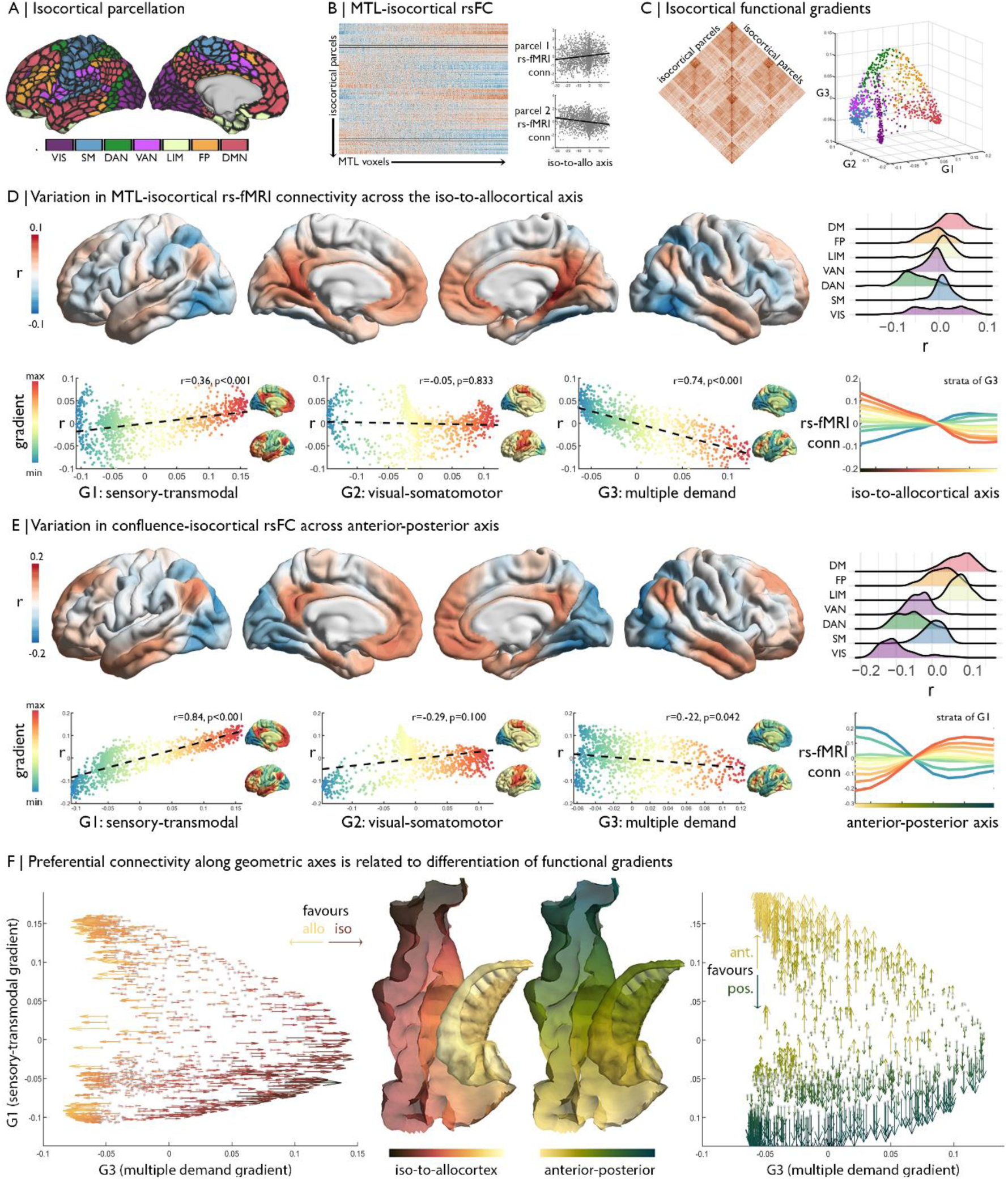
**A)** Parcellation of the isocortex, coloured by functional community ^64^. **B)** rs-fMRI connectivity between isocortical parcels and MTL voxels, with examples of row-wise associations with the iso-to-allocortical axis. **C)** Isocortical rs-fMRI connectivity was decomposed into a set of functional gradients, shown in a 3D space on the right. **D)** Surface maps show parcel-wise correlation of rs-fMRI connectivity with iso-to-allocortical axis. Red indicates increasing connectivity towards the allocortex, whereas blue indicates increasing functional connectivity towards the isocortex. The ridgeplot depicts the r values for each community. Scatter plots depict how regional associations with iso-to-allocortical axis (r, y-axis) vary as a function of the position each of the functional gradients (x-axis). Functional gradients are presented on the cortical surface insets right to each scatter plot. Line plot shows the average rs-fMRI connectivity ten strata of the third functional gradient with MTL voxels, organised along the iso-to-allocortical axis. **E)** We repeated the analysis in D for the anterior-posterior axis, showing a strong association with the sensory-transmodal gradient. **F)** Both scatterplots depict isocortical parcels in functional gradient space, defined by the sensory-transmodal and the multiple demand gradients. Arrows represent the correlation of rs-fMRI connectivity from that isocortical parcel with the iso-to-allocortical *(left)* and the anterior-posterior axes *(right)*. Arrow direction reflects the sign of the correlation, corresponding to the respective axis, and the colour depicts the strength.

The strength of rs-fMRI connectivity between the MTL and the isocortex varied as a function of the iso-to-allocortical axis. Specifically, lateral dorsal attention and fronto-parietal networks exhibited stronger rs-fMRI connectivity towards the isocortical anchor of the cortical confluence, whereas posterior cingulate and medial prefrontal regions of the default mode network and medial occipital areas in the visual network showed stronger connectivity towards the allocortical anchor (**Figure 3D**). A similar pattern was summarized by observing a parametric relation between position on the iso-to-allocortical axis and rs-fMRI along the third functional gradient (r=-0.74, p_spin_<0.001), which depicts a differentiation of the multiple demand system ^60,61^. The multiple demand anchor of the third functional gradient had stronger connectivity to isocortical compartments of the mesiotemporal confluence, whereas the opposing anchor comprising transmodal and sensory networks had higher connectivity with allocortical compartments. The association appears to be specific to the third functional gradient, supported by higher correlation coefficients than the first (z=12.30 p<0.0001) and second functional gradients (z=21.33 p<0.0001).

While we found that the iso-to-allocortical representation of the multiple demand gradient was preserved within thirds of the long-axis (correlation of r map with G3: 0.27<r<0.40), long axis position nevertheless affected representation of macroscale topographies specifically with respect to the first, sensory-transmodal functional gradient (correlation of r map with G1: -0.41<r<0.45). Further elucidating preferential rs-fMRI connectivity patterns of the anterior-posterior axis, we indeed found that this largely reflected connectivity transitions described by the sensory-transmodal functional gradient (r=-0.84, p_spin_<0.001; **Figure 3E**). The transmodal anchor had stronger functional connectivity with the anterior aspect of the MTL, while the sensory anchor had stronger connectivity with the posterior aspect.

Although our included datasets have overall high data quality, signal drop out in temporal lobe areas may potentially affect imaging findings. To dispel such concerns, we assessed signal to noise ratio (SNR) from the rs-fMRI data. While we indeed observed variations in both temporal as well as spatial SNR across the MTL, findings were virtually identical controlling for SNR via partial correlation [compared to the iso-to-allocortical axis from *Figure 3B*: r_spatial_=0.92, r_temporal_=0.99; compared to the anterior-posterior axis *from Figure 3C:* r_spatial_=0.99, r_temporal_=0.99]. Similarly, functional findings from mesiotemporal confluence were virtually consistent when excluding of temporal lobe regions as targets in the macrolevel functional connectivity findings (compared to *Figure 3F:* r=0.85).

Together, these analyses demonstrate an important interaction of the iso-to-allocortical and anterior-posterior axes of the MTL in understanding its functional relationship to the rest of the brain, with shifts in the iso-to-allo-cortical axis relating to connectivity to the multiple demand gradient and shifts along anterior-to-posterior axis relating to connectivity in the sensory-transmodal gradient **(Figure 3F)**.

### Replication of functional analyses

While the singular nature of BigBrain did not allow for additional cytoarchitectural cross-validations, we could test consistency of our *in vivo* functional analyses in an independent multi-modal imaging dataset (Human Connectome Project; see *Methods* for details). In line with the original analysis, we observed decreasing effective connectivity with increasing deviation from the iso-to-allocortical axis (r_iso_=-0.24; r_allo_=-0.47; **Figure 4A**), meaning there was stronger effective connectivity between closer neighbours on the iso-to-allocortical axis than to off axis nodes. Effects varied within anterior, middle and posterior thirds (**Figure 4A right**), again demonstrating intrinsic signal flow over iso-to-allocortical and anterior-posterior axes. Additionally, the iso-to-allocortical and anterior-posterior axes reflected distinct macroscale functional topographies. As in the original analyses, strength of isocortical rs-fMRI connectivity with the iso-to-allocortical axis corresponded with the multiple demand functional gradient (r=-0.52, p_spin_<0.001), whereas variations in rs-fMRI connectivity along the anterior-posterior axis corresponded with the sensory-transmodal functional gradient (r=0.33, p_spin_<0.001) (**Figure 4B**).

**Figure 4.**
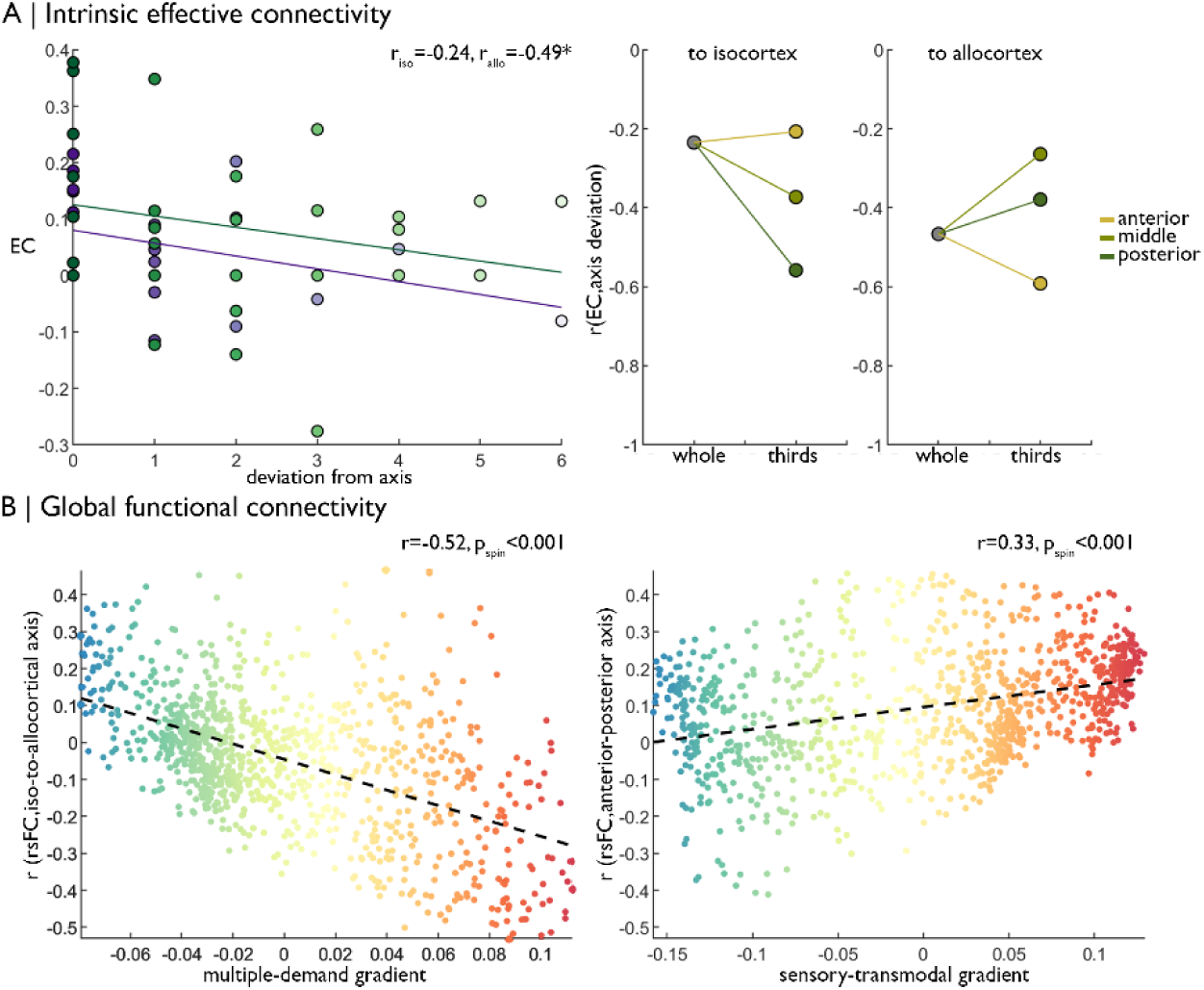
Replication of functional analyses in an independent dataset. **(A) *Left*** Strength of effective connectivity decreases with deviation from the iso-to-allocortical axis. Green = to isocortex. Purple = to allocortex. ***Right*** The correlation between effective connectivity and deviation from iso-to-allocortical axis differed within anterior, middle and posterior thirds. **(B)** Scatterplots show relationship between varying strength of rs-fMRI along the iso-to-allocortical *(left)* and anterior-posterior *(right)* axes mirror multiple demand and sensory-transmodal isocortical functional gradients, respectively.

## Discussion

Our study set out to determine whether macro and microstructural features of the human mesiotemporal lobe (MTL) offer an explanation for its broad influence on neural function. The MTL, a cytoarchitecturally unique region that harbours a transition from three-to-six layered cortex, has been a key focus of neuroscience for decades. Its intrinsic circuity is generally considered a model system to study how structural and functional properties co-vary in space ^65–67^. Furthermore, the MTL is known to have distributed connectivity patterns to multiple macroscale brain networks and to participate in numerous cognitive processes ^1–9,68^. Our work suggests that the intersection of the iso-to-allocortical cytoarchitectural gradient with the anterior-posterior axis allows the MTL to topographically represent multiple, dominant motifs of whole-brain functional organisation, which facilitates the distribution of neural codes computed within the internal hippocampal circuity across the cortical hierarchy.

Our starting point was an observer-independent characterisation of cytoarchitectural transitions in the MTL. To this end, we integrated manual hippocampal segmentation ^69^ with an isocortical surface model on an ultra-high resolution 3D histological reconstruction of the human brain ^62^. This approach allowed for a depth-wise microstructure profiling along a continuous geometric axis running through the folds of the MTL. Expanding from prior work focusing on specific MTL segments ^21,26,32^, several analyses specifically interrogated the transition from parahippocampal isocortex with a six-layered structure to three-layered hippocampal allocortex. A data-driven framework, based on unsupervised manifold learning of microstructure profile covariance ^70^, revealed a smooth and continuous iso-to-allocortical gradient as the principle axis of variation in cytoarchitecture in the MTL. Although our approach did not specifically aim at decomposing the MTL into discrete subfields with sharp boundaries (see ^71,72^ for histological and MRI efforts), the subfields of the MTL were nevertheless embedded within the iso-to-allocortical gradient and transitions from one subfield to the next were relatively smooth. One exception was CA2, however, which is recognized for high neuronal density ^69^ and unique synaptic properties within the larger CA complex ^73,74^.

Manifold learning was complemented by quantitative approaches to fingerprint cytoarchitecture based on statistical moments ^75–77^ and an evaluation of their accuracy in predicting location along the iso-to-allocortical axis for a given intracortical profile. Here, the most salient feature of the iso-to-allocortical transition was microstructure profile skewness, reflecting a relative increase in cell staining in uppermost surfaces. The uppermost molecular layer, sparsely populated by neurons in isocortex, appears less prominent towards the three-layered allocortex ^21^, an effect that is likely accentuated by decreasing cortical thickness ^21^. The pattern was conserved along the anterior-posterior axis, showing that the skewness-based cytoarchitectural gradient holds similar influence along the long axis. Finally, this iso-to-allocortical cytoarchitectural gradient extends upon the principle axis of microstructural differentiation identified across the isocortex ^76^. Together, this work shows a gradual differentiation in cyto-and myelo-architecture from sensory to limbic areas defines a core organizational axis in humans, extending from prior work in non-human primates and rodents ^70,76,78–81^.

As the next step, we assessed whether the cytoarchitectural gradient along the iso-to-allocortical axis confers the intrinsic functional organisation of the MTL, complementing previous modular descriptions of its internal circuity that posit dense connectivity between parahippocampal, perirhinal/entorhinal, and hippocampal subregions ^82,83^. Projecting the confluence model to high-definition *in vivo* MRI and studying estimates of directed connectivity, the evidence supported a dominant flow of connectivity along the iso-to-allocortical axis, in line with the purported link between cytoarchitectural similarity and inter-regional connectivity ^33,34,36,84,85^. Signal flow along iso-to-allocortical axis is known to be mediated by intracortical fiber tracts along the MTL infolding, notably mossy fibres and Schaefer collaterals ^38,86^. While signal flow and cytoarchitectural measurements are inextricably linked to distance, the diminishment of this pattern within anterior-posterior thirds of the MTL shows that it is not purely a product of spatial proximity. Following the relative preservation of iso-to-allocortical gradients along the MTL long axes, signal flow patterns were also statistically similar along different anterior-to-posterior segments. Yet, we nevertheless observed subtle differences, supporting interactions between anterior-posterior and iso-to-allocortical axes on the internal MTL circuitry. This effect is likely driven by the variable connectivity of the hippocampus to the surrounding isocortex along the anterior-posterior axis, rather than anterior-posterior variations in intrinsic connectivity of the hippocampus proper. Projections from the parahippocampus to the entorhinal cortex, subiculum and CA1 are clearly stratified into anterior (perirhinal) and posterior (postrhinal) tracts in the rodent brain ^87,88^, which however converge as early as the lateral entorhinal cortex ^89^. In line with this evidence, recent neuroimaging studies have shown more variability in rs-fMRI connectivity between anterior-posterior aspects of the MTL than between hippocampal subfields ^90^ and that subregions of the parahippocampal and entorhinal cortex exhibit preferential connectivity to either anterior or posterior aspects of the subiculum ^44^. Together, this evidence demonstrates that distinct anterior and posterior input streams to the hippocampus are integrated and flow along the iso-to-allocortical cytoarchitectural gradient. In other words, the steep descent in laminar differentiation across the cortical confluence underpins signal flow along the iso-to-allocortical axis and pulls together the anterior and posterior processing streams within the MTL.

We furthermore characterised the position of the MTL in the macroscale cortical landscape, using both modular and dimensional decompositions of whole-brain function ^56,57,64,91^. Particularly the latter framework has recently been employed to describe variations in microcircuitry ^79,92,93^, large-scale functional organisation ^18^, and putative neural hierarchies^20^. Our work demonstrated that the two major geometric axes of the MTL exhibited complementary affiliation to macroscale cortical systems and thereby balance distinct modes of cortical dynamics. The iso-to-allocortical axis is related to decreasing connectivity with multiple demand systems - multimodal areas involved in external attention across multiple contexts ^60,61^ - while the anterior-posterior axis reflected a shift from preference for transmodal cortex, such as the default mode network, to sensory-and-motor regions. Our results thus suggest that the anterior and posterior processing streams are successively integrated along steps of the iso-to-allocortical axis while external attentional processes are increasingly dampened. Higher-order representations constructed within the depths of the hippocampus then flow from allo-to-isocortex before being broadcast back to sensory and transmodal areas. This “hierarchy of associativity” is also evidenced by invasive tract-tracing in non-human animals ^24^ and could provide a neural workspace that is capable of representing both the abstract structure of tasks, as well as their specific sensory features, thereby supporting the hypothesised role of the MTL in cognitive maps ^15–17^. To our knowledge, no study has specifically examined continuous variations in functional connectivity along the iso-to-allocortical axis of the MTL; however, our results align with observed participation of the parahippocampus and fusiform gyri in externally-oriented and attention networks ^56,94,95^. The iso-to-allocortical axis also mirrors a reduction in the diversity of projections ^24^, which further explains the shift from involvement in large-scale polysynaptic networks toward more constrained, local connectivity. Sensory-transmodal differentiation of rs-fMRI connectivity along the MTL long axis is well-evidenced in the literature ^39,44,46,47,96^ and indicates more anatomical and topological proximity of anterior mesiotemporal components to default and fronto-parietal systems, while posterior hippocampal and parahippocampal regions are more closely implicated in sensory-spatial interactions with external contexts.

While we could replicate *in vivo* functional imaging findings across two independent datasets, among them the HCP benchmark repository, the singular nature of the BigBrain currently prohibits replication of our 3D histological findings. Follow up work based on ultra-high field magnetic resonance imaging may take similar approaches to explore MTL transitions with respect to *in vivo* cortical myeloarchitecture or intrinsic signal flow across cortical depths ^70,97–99^, thus allowing assessment of how inter-individual differences in structural and functional organisation of the MTL relate to inter-individual differences in cognitive and affective phenotypes. Given that structural and functional perturbations of the MTL are at the core of multiple brain diseases, notably Alzheimer’s disease, schizophrenia, and drug-resistant temporal lobe epilepsy ^100–105^, our framework may furthermore advance the understanding and management of these common and severe conditions. By making our MTL confluence model openly available [https://github.com/MICA-MNI/micaopen], we hope to facilitate such future investigations.

To conclude, our work shows that cytoarchitectural differentiation underpins patterns of intrinsic signal flow within the MTL microcircuit, while the long-axis allows for broad distribution of the computed neural code across the cortical hierarchy. By focusing on one of the most intriguing and integrative regions of the brain, this work illustrates how cytoarchitectural variation can underpin both the convergence of distributed processing streams and furthermore facilitate the distribution of integrated neural computations across large-scale systems. More broadly, our study of the MTL may provide clues for fundamental aspects of structure-function coupling in the brain ^18,24^ and in revealing core principles underlying macroscale cortical organisation ^22,28–30,106^.

## ACKNOWLEDGEMENTS

Casey Paquola was funded through a postdoctoral fellowship of the Fonds de la Recherche due Quebec – Santé (FRQ-S). Oualid Benkarim was funded by a Healthy Brains for Healthy Lives (HBHL) postdoctoral fellowship. Boris Bernhardt acknowledges research support from the National Science and Engineering Research Council of Canada (NSERC Discovery-1304413), the Canadian Institutes of Health Research (CIHR FDN-154298), SickKids Foundation (NI17-039), Azrieli Center for Autism Research (ACAR-TACC), and the Tier-2 Canada Research Chairs program. Andrea Bernasconi and Neda Bernasconi were funded by CIHR and NSERC. Jessica Royer was funded by a CIHR fellowship. Jonathan Smallwood was funded by the European Research Council (WANDERINGMINDS). Sara Lariviere acknowledges funding from Fonds de la Recherche du Québec – Santé (FRQ-S) and the Canadian Institutes of Health Research (CIHR). We would also like to acknowledge support from the Helmholtz Foundation and the Healthy Brains for Healthy Lives initiative.

## Methods

### Histological model of cortical confluence

An ultra-high resolution Merker stained 3D volumetric histological reconstruction of a *post mortem* human brain from a 65-year-old male was obtained from the open-access BigBrain repository, along with pial and white matter surface reconstructions ^62^. We also obtained manually segmented hippocampal subregions, which were labelled with a three-way internal coordinate system; anterior-posterior, proximal-distal and inner-outer based on the solving Laplace’s equation (DeKraker et al., 2019; https://osf.io/x542s/). We generated continuous inner and outer surfaces of the hippocampus by triangulating coordinates at the minimum and the maximum of an inner-outer axis ^69^. The inner surface of the hippocampus, with respect to the hippocampal fissure, is continuous with the pial surface, whereas the outer surface of the hippocampus is continuous with the white matter boundary ^38^. Next, we designated the most proximal vertices as hippocampal bridgeheads (*i*.*e*., closest to isocortex) and generated links via nearest neighbour interpolation to the entorhinal, parahippocampal or fusiform isocortex vertices, with an additional constraint that the nearest neighbour is inferior to the bridgehead in order to follow the curvature of the MTL. This procedure was developed and evaluated based on visual inspection by a trained anatomist (CP). The entorhinal, parahippocampal, and fusiform areas were defined using the Desikan-Killany atlas, nonlinearly transformed to the BigBrain histological surfaces ^107^, and the hippocampal subfields were assigned in line with manual segmentation ^69^. Only the right hemisphere was investigated, owing to a tear in the left entorhinal cortex in BigBrain. We remapped the proximal-distal axis as the minimum geodesic distance to a hippocampal bridgehead to ensure equal scaling on either side of the confluence, and hereafter use this metric as the iso-to-allocortical axis. Negative values represent vertices before the hippocampal bridgehead, and positive values are within the hippocampus. The y-coordinate was used to capture the anterior-posterior axis of the whole MTL.

### Cytoarchitectural mapping of the cortical confluence

We generated 14 equivolumetric surfaces between the inner and outer confluent surfaces and sampled the intensities from the 40μm BigBrain blocks along 9432 matched vertices, creating microstructure profiles in the direction of cortical columns for the hippocampus and adjacent MTL areas. Staining intensities reflect cellular density and soma size ^62,108^. In lieu of precise laminar decomposition, which is currently untenable in such transition regions ^109^, we utilised an equivolumetric algorithm was used to optimise the fit of depth-wise surfaces with variations in thickness and curvature ^110^. For the sake of clarity, we refer to the surfaces by the percentage depth of their initialisation, which was adjusted for curvature to become equivolumetric.

Unsupervised machine learning tested the hypothesis that the iso-to-allocortical axis is the principle gradient of cytoarchitectural differentiation in the MTL. The procedure involved calculating microstructure profile covariance between all vertices then extracting the principle axis of cytoarchitectural variation as the first eigenvector from diffusion map embedding ^70,91,111^. Diffusion map embedding is optimally suited for multi-scale architectures, because unlike global methods, such as principle component analysis and multidimensional scaling, local geometries are preserved and then integrated into a set of global eigenvectors. The statistical relationship between the principle eigenvector and the proximal-distal axis was modelled with a series of increasingly complex polynomial curves. We selected the model at the inflection point of adjusted R^2^ values, taken across polynomials of 1-5. We extracted the residuals from the selected model to determine which regions deviated from the expected association.

The defining cytoarchitectural features of the iso-to-allocortical axis were established using feature selection in a random forest regression. A set of 20 cytoarchitectural features [16 depth-wise intensities and 4 central moments ^76^] were z-standardised and fed into random forest regression to predict iso-to-allocortical axis. Hyperparameters were selected via 5-fold cross validation in a grid search. We trained the model on 70% of vertices then measured prediction accuracy by R^2^ with the remaining 30%. Feature importance was estimated as the mean decrease in variance, and features with higher than average importance were selected. The procedure was repeated across 100 splits to assess robustness of prediction accuracy and feature importance. Using the same hyperparameters, we refit the model with only the selected features and assessed the predictive accuracy in the same manner. For post-hoc inference, we assessed the pattern of change in each selected feature with the iso-to-allocortical axis using a cubic polynomial (the best fit in the unsupervised model; **Figure 1F**).

To assess the consistency of iso-to-allocortical variations along the anterior-posterior axis, we repeated our analyses with stratification by the anterior-posterior axis. We fit a cubic polynomial between each selected feature and the iso-to-allocortical axis within 1mm coronal slices of confluence model and approximated the goodness of fit (adjusted R^2^) for the 23 coronal slices that included the hippocampus proper.

### Registration to MNI152 standard atlas

The cortical confluence model was non-linearly registered to the MNI152 2mm standard atlas to facilitate targeted associations to *in vivo* functional imaging. This procedure involved labelling voxels in native histological space (1000μm) according to their position along the iso-to-allocortical axis. To allow for downsampling (40→1000μm), we used average axis values of all vertices contained within a voxel. Next, we applied a series of one linear and two nonlinear transformations to register the volumetric representation of cortical confluence to MNI152 standard atlas. The first two transformations are available via the BigBrain repository (ftp://bigbrain.loris.ca/BigBrainRelease.2015/3D_Volumes/MNI-ICBM152_Space/), and the final nonlinear optimisation is available as part of an open science framework (Xiao et al., 2019; https://osf.io/xkqb3/). Finally, we restricted the model to the anterior-posterior range of the hippocampus proper to allow appropriate comparisons of both geometric axes and used a grey matter partial volume mask to ensure the standardised volumetric atlas conformed to cortical boundaries.

### Functional MRI acquisition and preprocessing

We studied resting state functional imaging data from 40 healthy adults (microstructure informed connectomics (MICs) cohort; 14 females, mean±SD age=30.4±6.7, 2 left-handed). Participants gave informed consent and the study was approved by the local Research Ethics Board of the Montreal Neurological Institute and Hospital. MRI data was acquired on a 3T Siemens Magnetom Prisma-Fit with a 64-channel head coil. A submillimetric T1-weighted image was acquired using a 3D-MPRAGE sequence (0.8mm isotropic voxels, 320×320 matrix, 24 sagittal slices, TR=2300ms, TE=3.14ms, TI=900ms, flip angle=9°, iPAT=2). One 7 min rs-fMRI scan was acquired using multi-band accelerated 2D-BOLD echo-planar imaging (TR=600ms, TE=30ms, 3mm isotropic voxels, flip angle=52°, FOV=240×240mm^2^, slice thickness=3mm, mb factor=6, echo spacing=0.54mms). Participants were instructed to keep their eyes open, look at fixation cross, and not fall asleep. All fMRI data underwent gradient unwarping, motion correction, distortion correction, brain-boundary-based registration to structural T1-weighted scan and grand-mean intensity normalization. The rs-fMRI data was additionally denoised using an in-house trained ICA-FIX classifier ^113,114^, which notably outperforms motion scrubbing and spike regression in reproducing resting state networks and conservation of temporal degrees of freedom ^115,116^. Timeseries were sampled on native cortical surfaces and averaged within 1000 spatially contiguous, functionally defined parcels (Schaefer et al., 2018). Finally, for each subject, we generated a matrix reflecting a two-step linear transformation (MNI152 to native T1-weigthed to native rs-fMRI space) to facilitate mapping the standardised volumetric atlas to individual subjects’ functional images.

### Dynamic modelling

Spectral dynamic causal modelling estimated the effective connectivity between regions of the cortical confluence model ^63,117^. This approach involves a generative model with one state depicting the interactions of neuronal populations and another state providing a conventional linear convolution of neuronal activity to the hemodynamic response ^118^. We specified a fully connected dynamic model, without exogenous inputs, and estimated model parameters, that is the effective connectivity between regions, and their marginal likelihood using a Variational Laplace inversion ^119^. In line with previous research ^63,117^, we used a fourth order autoregressive model to estimate cross spectra of the timeseries. The model was built for each of the 40 subjects separately, and group-level estimates were calculated using a second-level parametric empirical Bayesian model ^120^. The standardised volumetric atlas of the iso-to-allocortical axis was divided into a set of discrete bins and transformed to individual functional space, then BOLD timeseries were extracted and the mean timeseries was calculated for each bin. We evaluated the optimal number of bins, ranging from 4 and 14, and selected the finest resolution that still yielded a hierarchical model for which inversion was computationally feasible using a second-level parametric empirical Bayesian analysis. The percentage of voxels of each bin that occupied anatomical subregions was calculated based on the Desikan-Killany atlas and hippocampal subfield segmentations in FreeSurfer ^121,122^. Next, we applied Bayesian model reduction to the posterior densities over the second level parameters, involving a greedy search over all permutations of parameter inclusion, to discover the structure of the optimal sparse graph.

To test the hypothesis that the iso-to-allocortical axis constrains the signal flow through the mesiotemporal confluence, we compared the strength of effectivity connectivity with the concordance of an edge to the iso-to-allocortical axis using Pearson correlation. Each edge was labelled by the degree of deviation from the iso-to-allocortical axis (0-6 in the 8-bin model). We further stratified these edges by the direction along the axis (*i*.*e*., signal flow towards isocortex or towards allocortex). Next, we divided the standardised volumetric atlas into thirds along the anterior-posterior axis and repeated the dynamic model with the preselected number of bins for each third separately. We assessed consistency across the anterior-posterior axis by calculating the Pearson correlation coefficient of the effective connectivity estimates from non-reduced models between y-axis restricted and the full mesiotemporal confluence model, as well as repeating the association of the effective connectivity with deviation from the iso-to-allocortical axis in the reduced model obtained from Bayesian model reduction.

### Extrinsic functional connectivity

The association between iso-to-allocortical axis and macroscale rs-fMRI connectivity patterns was estimated via a Pearson correlation for each isocortical parcel. Positive r values reflect higher connectivity towards the allocortical end, whereas negative r values reflect higher connectivity towards the isocortical end. We compared the spatial pattern of r values across the isocortex with canonical functional gradients ^57^. To this end, we calculated parcel-parcel rs-fMRI connectivity matrix for each participant, averaged across the cohort, transformed this into a normalised angle matrix, and subjected it to diffusion map embedding. The principle eigenvectors, referred to as functional gradients, resembled previous descriptions ^57,70,123^. Spearman correlations assessed the spatial correspondence of the r map with the functional gradients and spin permutations established significance, while accounting for spatial autocorrelation ^124,125^. We tested the specificity of the associations with functional gradients by comparing the correlation coefficients ^126^. We repeated this comparison within thirds of the anterior-posterior axis and tested the association of rs-fMRI connectivity to the isocortex with the anterior-posterior axis.

### Effects of SNR

We assessed potential effects of SNR variations on macroscale MTL connectivity. Here, spatial SNR was taken as the mean signal within a voxel divided by the standard deviation of the signal across all grey matter voxels. Temporal SNR was taken as the mean signal of a voxel divided by the standard deviation of signal in that voxel over time. Additionally, we assessed the contribution of the temporal lobe to the relationship between spatial maps and functional gradients by excluding the lobe for computing the final correlation. We used the Deskian-Killany atlas to define the temporal lobe and excluded isocortical parcel with any vertices in the temporal lobe (n=196).

### Replication

We replicated the fMRI findings in 40 unrelated healthy adults (14 females, age=29.4±3.8 years) from the minimally preprocessed S900 release of the Human Connectome Project (HCP) ^127^.

#### a) MRI acquisition

In brief, MRI data were acquired on the HCP’s custom 3T Siemens Skyra equipped with a 32-channel head coil. Two T1-weighted images with identical parameters were acquired using a 3D-MPRAGE sequence (0.7 mm isotropic voxels, matrix = 320 × 320, 256 sagittal slices; TR = 2,400 ms, TE = 2.14 ms, TI = 1,000 ms, flip angle = 8°; iPAT = 2). Four rs-fMRI scans were acquired using multiband accelerated 2D-BOLD echo-planar imaging (2 mm isotropic voxels, matrix = 104 × 90, 72 sagittal slices; TR = 720 ms, TE = 33 ms, flip angle = 52°; mb factor = 8; 1,200 volumes/scan). We utilised the first session of rs-fMRI. Participants were instructed to keep their eyes open, look at fixation cross, and not fall asleep. For rs-fMRI, the timeseries were corrected for gradient nonlinearity and head motion.

#### b) MRI processing

The R-L/L-R blipped scan pairs were used to correct for geometric distortions. Distortion-corrected images were warped to T1w space using a combination of rigid body and boundary-based linear registrations ^128^. These transformations were concatenated with the transformation from native T1w to MNI152 to warp functional images to MNI152. Further processing removed the bias field (as calculated for the structural image), extracted the brain, and normalised whole-brain intensity. A high-pass filter (>2,000s FWHM) corrected the timeseries for scanner drifts, and additional noise was removed using ICA-FIX ^113^. Tissue-specific signal regression was not performed ^129^. Using the functional images warped to MNI152 space, timeseries were extracted from all voxels encompassed by the cortical confluence atlas and 1000 isocortical parcels of the Schaefer 7-network atlas ^64^. Timeseries were averaged within each isocortical parcel.

#### c) fMRI analysis

To analyze intrinsic circuitry, we tested whether effective connectivity was stronger on the iso-to-allocortical axis, using an 8bin iso-to-allocortical division of the entire MTL. We repeated this analysis within three consecutive segments of the long axis. To analyze macroscale connectivity, we calculated Pearson correlations between connectivity patterns of specific iso-to-allocortical MTL subsections and functional gradients. The isocortical functional gradients were re-generated within the HCP cohort and were highly similar to the original dataset.

## Supplementary Figures

**Figure S1:**
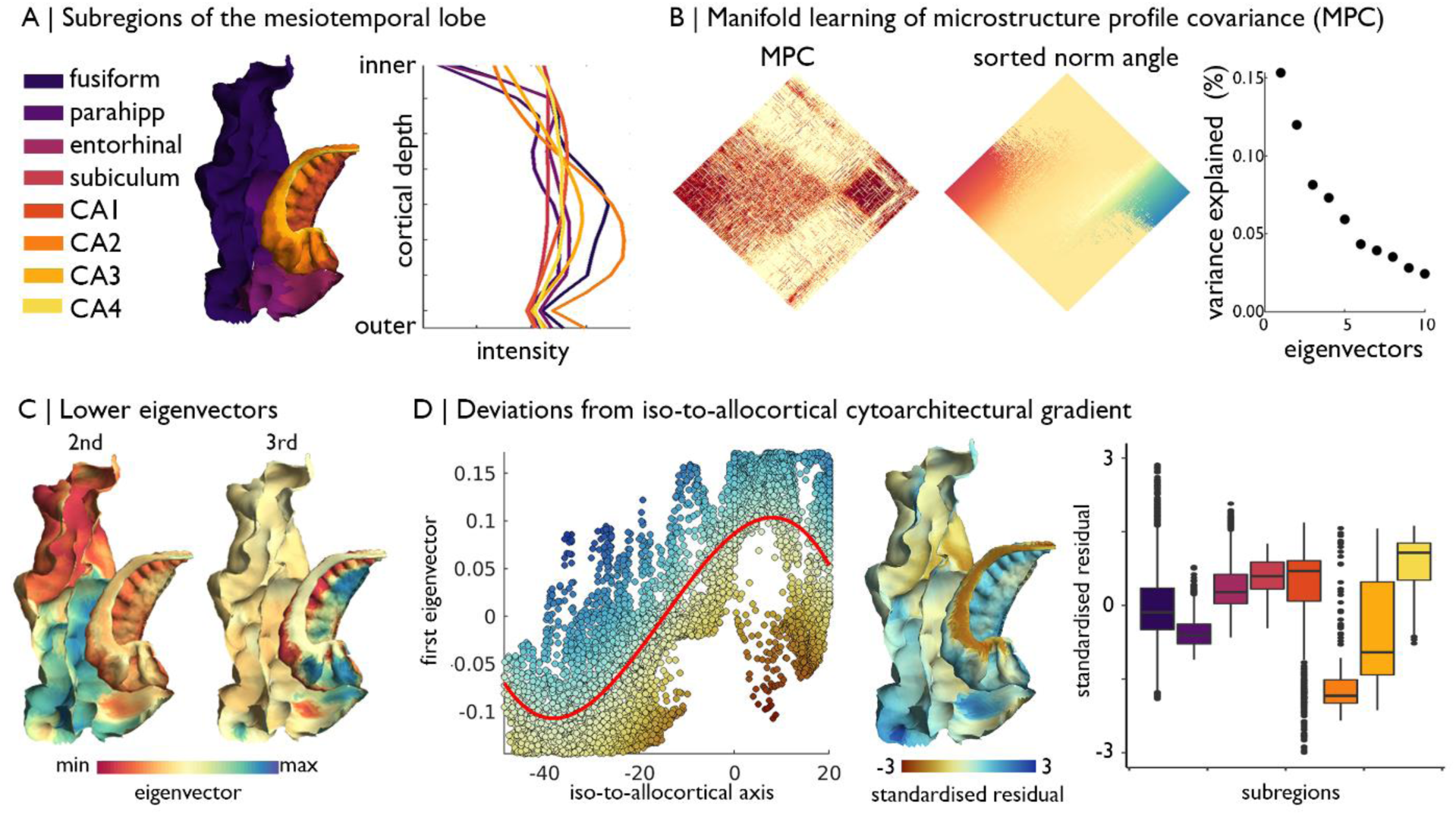
**(A)** Surface projection and average microstructure profiles of MTL subregions, defined by the Desikan-Killany atlas for isocortex and manual segmentations for the hippocampus proper. **(B)** Microstructure profile covariance (MPC) across all vertices in the cortical confluence model was transformed into a normalised angle matrix and sorted according to the first eigenvector from diffusion map embedding. Scatterplot shows approximate variance explained by each eigenvector. **(C)** Second and third eigenvectors shown on the MTL confluence model. **(D)** Deviations from the polynomial association of the geometric iso-to-allocortical axis with the first eigenvector were estimated by the standardised residuals. Standardised residuals are shown on the surface projection and stratified by subregions.

**Figure S2:**
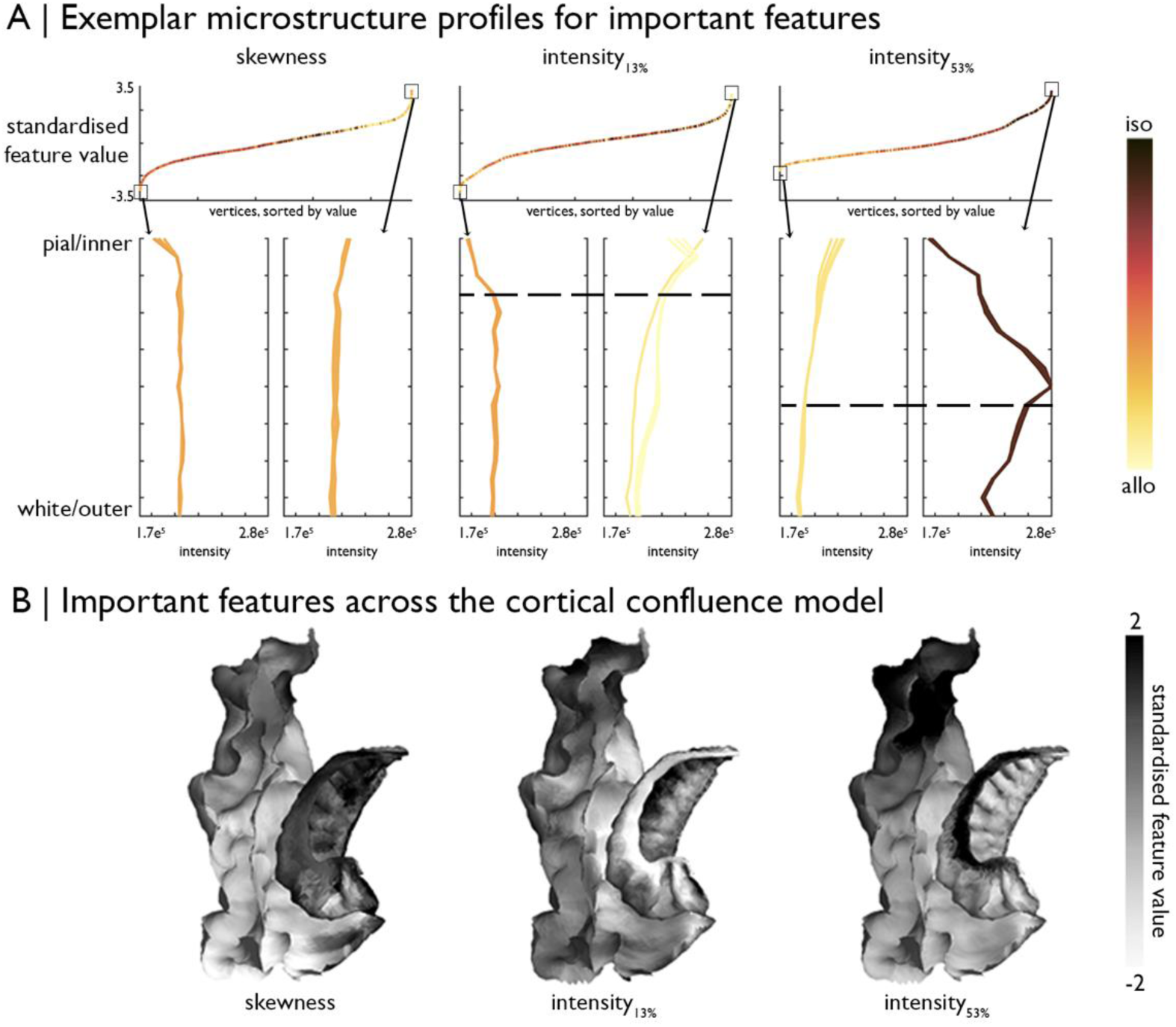
Exemplar microstructure profiles of selected features, namely skewness, intensity at ∼13% depth and ∼53% depth. **A)** *Top:* Standardised feature values of each vertex, sorted by value and coloured by position on the iso-to-allocortical axis. *Bottom:* Profiles of vertices with five highest and five lowest values of each feature. Again, vertices are coloured by position on the iso-to-allocortical axis. Higher Merker-staining intensity values reflect higher cellular density/soma size. **B)** Feature values of each vertex projected onto the MTL confluence model.

**Figure S3:**
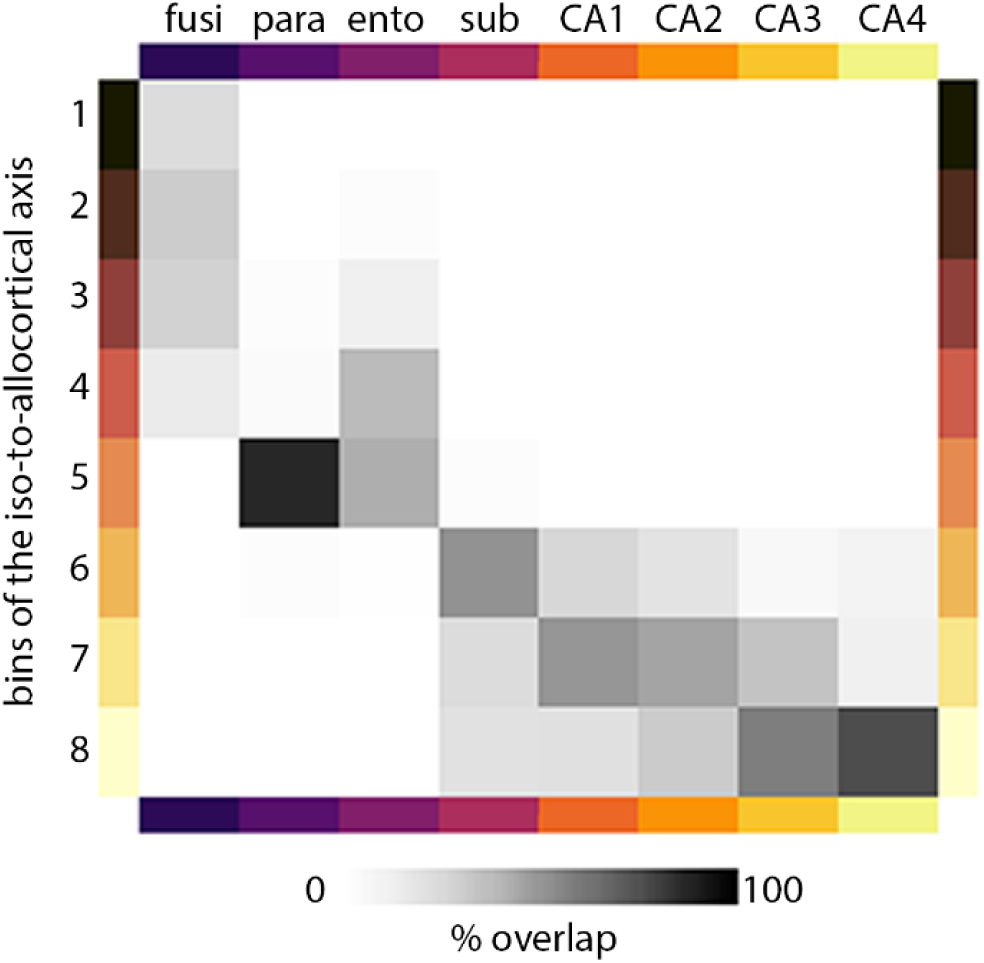
Characteristics of bins used in the dynamic modeling analysis. Percentage overlap of bins with subregions.

## Notes

### Competing Interest Statement

The authors have declared no competing interest.

## References

1. Moscovitch, M. et al. Functional neuroanatomy of remote episodic, semantic and spatial memory: A unified account based on multiple trace theory. Journal of Anatomy 207, 35–66 (2005).

2. Squire, L. R., Stark, C. E. L. & Clark, R. E. The Medial Temporal Lobe. Annu. Rev. Neurosci. 27, 279–306 (2004).

3. Milner, B. The medial temporal-lobe amnesic syndrome. Psychiatric Clinics of North America 28, 599–611 (2005).

4. Eichenbaum, H., Yonelinas, A. P. & Ranganath, C. The Medial Temporal Lobe and Recognition Memory. Annu. Rev. Neurosci. 30, 123–152 (2007).

5. Wang, D. et al. Comprehensive functional genomic resource and integrative model for the human brain. Science 362, eaat8464 (2018).

6. Eacott, M. J., Gaffan, D. & Murray, E. A. Preserved Recognition Memory for Small Sets, and Impaired Stimulus Identification for Large Sets, Following Rhinal Cortex Ablations in Monkeys. Eur. J. Neurosci. 6, 1466–1478 (1994).

7. Lee, A. C. H. & Rudebeck, S. R. Human medial temporal lobe damage can disrupt the perception of single objects. J. Neurosci. 30, 6588–6594 (2010).

8. Buzsáki, G. & Moser, E. I. Memory, navigation and theta rhythm in the hippocampal-entorhinal system. Nature Neuroscience 16, 130–138 (2013).

9. Felix-Ortiz, A. C. & Tye, K. M. Amygdala inputs to the ventral hippocampus bidirectionally modulate social behavior. J. Neurosci. 34, 586–595 (2014).

10. Lech, R. K. & Suchan, B. The medial temporal lobe: Memory and beyond. Behavioural Brain Research 254, 45–49 (2013).

11. Zheng, J. et al. Amygdala-hippocampal dynamics during salient information processing. Nat. Commun. 8, 1–11 (2017).

12. Saksida, L. M. & Bussey, T. J. The representational-hierarchical view of amnesia: Translation from animal to human. Neuropsychologia 48, 2370–2384 (2010).

13. Staresina, B. P. & Davachi, L. Mind the Gap: Binding Experiences across Space and Time in the Human Hippocampus. Neuron 63, 267–276 (2009).

14. Diana, R. A., Yonelinas, A. P. & Ranganath, C. Imaging recollection and familiarity in the medial temporal lobe: a three-component model. Trends Cogn. Sci. 11, 379–386 (2007).

15. O’Keefe, J. & Nadel, L. The hippocampus as a cognitive map. 570 (1978).

16. Tolman, E. C. Cognitive maps in rats and men. Psychol. Rev. 55, 189–208 (1948).

17. Behrens, T. E. J. et al. What Is a Cognitive Map? Organizing Knowledge for Flexible Behavior. Neuron 100, 490–509 (2018).

18. Mesulam, M.-M. From sensation to cognition. Brain 121, 1013–1052 (1998).

19. Felleman, D. J. & Van Essen, D. C. Distributed hierarchical processing in the primate cerebral cortex. Cereb. Cortex 1, 1–47 (1991).

20. Chanes, L. & Barrett, L. F. Redefining the Role of Limbic Areas in Cortical Processing. Trends Cogn. Sci. 20, 96–106 (2016).

21. Insausti, R., Muñoz-López, M., Insausti, A. M. & Artacho-Pérula, E. The human periallocortex: Layer pattern in presubiculum, parasubiculum and entorhinal cortex. A review. Front. Neuroanat. 11, (2017).

22. Sanides, F. Die Architektonik des menschlichen Stirnhirns zugleich eine Darstellung der Prinzipien seiner Gestaltung als Spiegel der stammgeschichtlichen Differenzierung der Grosshirnrinde. 201 (Springer, 1962).

23. Vogt, C. & Vogt, O. Allgemeinere Ergebnisse unserer Hirnforschung. J. für Psychol. und Neurol. 25, 279–461 (1919).

24. Lavenex, P. & Amaral, D. G. Hippocampal-neocortical interaction: A hierarchy of associativity. Hippocampus 10, 420–430 (2000).

25. García-Cabezas, M., Joyce, M. K. P., John, Y. J., Zikopoulos, B. & Barbas, H. Mirror trends of plasticity and stability indicators in primate prefrontal cortex. Eur. J. Neurosci. 46, 2392–2405 (2017).

26. Duvernoy, H. M., Cattin, F., Risold, P. Y., Vannson, J. L. & Gaudron, M. *The human hippocampus: Functional anatomy, vascularization and serial sections with MRI, fourth edition*. The Human Hippocampus: Functional Anatomy, Vascularization and Serial Sections with MRI, Fourth Edition (Springer Berlin Heidelberg, 2013). doi:10.1007/978-3-642-33603-4

27. Braak, H. & Braak, E. On areas of transition between entorhinal allocortex and temporal isocortex in the human brain. Normal morphology and lamina-specific pathology in Alzheimer’s disease. Acta Neuropathol. 68, 325–332 (1985).

28. Dart, R. A. The dual structure of the neopallium: Its history and significance. J. Anat. 69, 3 (1934).

29. Abbie, A. A. The excitable cortex in the Monotremata. (Australian Journal of Experimental Biology and Medical Science, 1938).

30. Abbie, A. A. Cortical lamination in a polyprotodont marsupial, Perameles nasuta. J. Comp. Neurol. 76, 509–536 (1942).

31. Sanides, F. Comparative Archiectonics of the Neocortex of Mammals and their Evolutionary Interpretation. Ann. N. Y. Acad. Sci. 167, 404–423 (1969).

32. Insausti, R. & Amaral, D. G. Hippocampal Formation. in The Human Nervous System 896–942 (Elsevier Inc., 2012). doi:10.1016/B978-0-12-374236-0.10024-0

33. Barbas, H. Pattern in the laminar origin of corticocortical connections. J. Comp. Neurol. 252, 415–422 (1986).

34. Hilgetag, C. C. & Grant, S. Cytoarchitectural differences are a key determinant of laminar projection origins in the visual cortex. Neuroimage 51, 1006–1017 (2010).

35. Beul, S. F., Grant, S. & Hilgetag, C. C. A predictive model of the cat cortical connectome based on cytoarchitecture and distance. Brain Struct. Funct. 220, 3167–3184 (2015).

36. Beul, S. F., Barbas, H. & Hilgetag, C. C. A Predictive Structural Model of the Primate Connectome. Sci. Rep. 7, (2017).

37. García-Cabezas, M. Á., Zikopoulos, B. & Barbas, H. The Structural Model: a theory linking connections, plasticity, pathology, development and evolution of the cerebral cortex. Brain Struct. Funct. 1–24 (2019). doi:10.1007/s00429-019-01841-9

38. Amaral, D. G. & Witter, M. P. The three-dimensional organization of the hippocampal formation: A review of anatomical data. Neuroscience 31, 571–591 (1989).

39. Strange, B. A., Witter, M. P., Lein, E. S. & Moser, E. I. Functional organization of the hippocampal longitudinal axis. Nature Reviews Neuroscience 15, 655–669 (2014).

40. Witter, M. P., Wouterlood, F. G., Naber, P. A. & Van Haeften, T. Anatomical Organization of the Parahippocampal-Hippocampal Network. Ann. N. Y. Acad. Sci. 911, 1–24 (2006).

41. Poppenk, J., Evensmoen, H. R., Moscovitch, M. & Nadel, L. Long-axis specialization of the human hippocampus. Trends Cogn. Sci. 17, 230–240 (2013).

42. Vogel, J. W. et al. A molecular gradient along the longitudinal axis of the human hippocampus informs large-scale behavioral systems. Nat. Commun. 11, 1–17 (2020).

43. Navarro Schröder, T., Haak, K. V., Jimenez, N. I. Z., Beckmann, C. F. & Doeller, C. F. Functional topography of the human entorhinal cortex. Elife 4, 1–17 (2015).

44. Maass, A., Berron, D., Libby, L. A., Ranganath, C. & Düzel, E. Functional subregions of the human entorhinal cortex. Elife 4, 1–20 (2015).

45. Zhong, Q. et al. Functional parcellation of the hippocampus from resting-state dynamic functional connectivity. Brain Res. 1715, 165–175 (2019).

46. Vos de Wael, R. et al. Anatomical and microstructural determinants of hippocampal subfield functional connectome embedding. Proc. Natl. Acad. Sci. U. S. A. 115, 10154–10159 (2018).

47. Przeździk, I., Faber, M., Fernández, G., Beckmann, C. F. & Haak, K. V. The functional organisation of the hippocampus along its long axis is gradual and predicts recollection. Cortex 119, 324–335 (2019).

48. Smallwood, J. et al. Representing representation: Integration between the temporal lobe and the posterior cingulate influences the content and form of spontaneous thought. PLoS One 11, (2016).

49. Vincent, J. L. et al. Coherent spontaneous activity identifies a hippocampal-parietal memory network. J. Neurophysiol. 96, 3517–3531 (2006).

50. Ojemann, J. G. et al. Anatomic localization and quantitative analysis of gradient refocused echo-planar fMRI susceptibility artifacts. Neuroimage 6, 156–167 (1997).

51. Braga, R. M. & Buckner, R. L. Parallel Interdigitated Distributed Networks within the Individual Estimated by Intrinsic Functional Connectivity. Neuron 95, 457-471.e5 (2017).

52. Rolls, E. T. Pattern separation, completion, and categorisation in the hippocampus and neocortex. Neurobiol. Learn. Mem. 129, 4–28 (2016).

53. Neunuebel, J. P. & Knierim, J. J. CA3 retrieves coherent representations from degraded input: Direct evidence for CA3 pattern completion and dentate gyrus pattern separation. Neuron 81, 416–427 (2014).

54. Staresina, B. P. et al. Hippocampal pattern completion is linked to gamma power increases and alpha power decreases during recollection. Elife 5, (2016).

55. Nakazawa, K. et al. Requirement for hippocampal CA3 NMDA receptors in associative memory recall. Science (80-.). 297, 211–218 (2002).

56. Yeo, B. T. T. et al. The organization of the human cerebral cortex estimated by intrinsic functional connectivity. J. Neurophysiol. 106, 1125–1165 (2011).

57. Margulies, D. S. et al. Situating the default-mode network along a principal gradient of macroscale cortical organization. Proc. Natl. Acad. Sci. 113, 12574–12579 (2016).

58. Murphy, C. et al. Distant from input: Evidence of regions within the default mode network supporting perceptually-decoupled and conceptually-guided cognition. Neuroimage 171, 393–401 (2018).

59. Murphy, C. et al. Modes of operation: A topographic neural gradient supporting stimulus dependent and independent cognition. Neuroimage 186, 487–496 (2019).

60. Fedorenko, E., Duncan, J. & Kanwisher, N. Broad domain generality in focal regions of frontal and parietal cortex. Proc. Natl. Acad. Sci. U. S. A. 110, 16616–16621 (2013).

61. Duncan, J. The multiple-demand (MD) system of the primate brain: mental programs for intelligent behaviour. Trends in Cognitive Sciences 14, 172–179 (2010).

62. Amunts, K. et al. BigBrain: An Ultrahigh-Resolution 3D Human Brain Model. Science (80-.). 340, 1472–1475 (2013).

63. Friston, K. J., Kahan, J., Biswal, B. & Razi, A. A DCM for resting state fMRI. Neuroimage 94, 396–407 (2014).

64. Schaefer, A. et al. Local-Global Parcellation of the Human Cerebral Cortex from Intrinsic Functional Connectivity MRI. Cereb. Cortex 28, 3095–3114 (2018).

65. Cembrowski, M. S. et al. Dissociable Structural and Functional Hippocampal Outputs via Distinct Subiculum Cell Classes. Cell 173, 1280-1292.e18 (2018).

66. Cembrowski, M. S., Wang, L., Sugino, K., Shields, B. C. & Spruston, N. Hipposeq: A comprehensive RNA-seq database of gene expression in hippocampal principal neurons. Elife 5, (2016).

67. Masukawa, L. M., Benardo, L. S. & Prince, D. A. Variations in electrophysiological properties of hippocampal neurons in different subfields. Brain Res. 242, 341–344 (1982).

68. Wang, S. F., Ritchey, M., Libby, L. A. & Ranganath, C. Functional connectivity based parcellation of the human medial temporal lobe. Neurobiol. Learn. Mem. 134, 123–134 (2016).

69. DeKraker, J., Lau, J. C., Ferko, K. M., Khan, A. R. & Köhler, S. Hippocampal subfields revealed through unfolding and unsupervised clustering of laminar and morphological features in 3D BigBrain. Neuroimage 116328 (2019). doi:10.1016/j.neuroimage.2019.116328

70. Paquola, C. et al. Microstructural and functional gradients are increasingly dissociated in transmodal cortices. PLOS Biol. 17, e3000284 (2019).

71. de Flores, R. et al. Characterization of hippocampal subfields using ex vivo MRI and histology data: Lessons for in vivo segmentation. Hippocampus (2019). doi:10.1002/hipo.23172

72. Wisse, L. E. M. et al. A harmonized segmentation protocol for hippocampal and parahippocampal subregions: Why do we need one and what are the key goals? Hippocampus 27, 3–11 (2017).

73. Dudek, S. M., Alexander, G. M. & Farris, S. Rediscovering area CA2: Unique properties and functions. Nature Reviews Neuroscience 17, 89–102 (2016).

74. Lorente de Nó, R. Studies on the structure of the cerebral cortex. II. Continuation of the study of the ammonic system. J. für Psychol. und Neurol. 46, 113–177 (1934).

75. Zilles, K., Schleicher, A., Palomero-Gallagher, N. & Amunts, K. Quantitative Analysis of Cyto- and Receptor Architecture of the Human Brain. in Brain Mapping: The Methods 573–602 (Academic Press, 2002). doi:10.1016/b978-012693019-1/50023-x

76. Paquola, C. et al. Shifts in myeloarchitecture characterise adolescent development of cortical gradients. Elife 8, (2019).

77. Schleicher, A., Amunts, K., Geyer, S., Morosan, P. & Zilles, K. Observer-Independent Method for Microstructural Parcellation of Cerebral Cortex: A Quantitative Approach to Cytoarchitectonics. Neuroimage 9, 165–177 (1999).

78. Burt, J. B. et al. Hierarchy of transcriptomic specialization across human cortex captured by structural neuroimaging topography. Nat. Neurosci. 21, 1251–1259 (2018).

79. Fulcher, B. D., Murray, J. D., Zerbi, V. & Wang, X.-J. Multimodal gradients across mouse cortex. Proc. Natl. Acad. Sci. 116, 4689–4695 (2019).

80. Huntenburg, J. M., Bazin, P.-L. & Margulies, D. S. Large-Scale Gradients in Human Cortical Organization. Trends Cogn. Sci. 22, 21–31 (2018).

81. Huntenburg, J. M. et al. A Systematic Relationship Between Functional Connectivity and Intracortical Myelin in the Human Cerebral Cortex. Cereb. Cortex 27, 981–997 (2017).

82. Lacy, J. W. & Stark, C. E. L. Intrinsic functional connectivity of the human medial temporal lobe suggests a distinction between adjacent MTL cortices and hippocampus. Hippocampus 22, 2290–2302 (2012).

83. Shah, P. et al. Mapping the structural and functional network architecture of the medial temporal lobe using 7T MRI. Hum. Brain Mapp. 39, 851–865 (2018).

84. Barbas, H. & Pandya, D. N. Architecture and intrinsic connections of the prefrontal cortex in the rhesus monkey. J. Comp. Neurol. 286, 353–375 (1989).

85. Barbas, H. & Rempel-Clower, N. Cortical structure predicts the pattern of corticocortical connections. Cereb. Cortex 7, 635–646 (1997).

86. Zeineh, M. et al. Direct Visualization and Mapping of the Spatial Course of Fiber Tracts at Microscopic Resolution in the Human Hippocampus. Cereb. Cortex 27, 1779–1794 (2017).

87. Naber, P. A., Witter, M. P. & Da Silva, F. H. L. Perirhinal cortex input to the hippocampus in the rat: evidence for parallel pathways, both direct and indirect. A combined physiological and anatomical study. Eur. J. Neurosci. 11, 4119–4133 (1999).

88. Naber, P. A., Witter, M. P. & Lopes da Silva, F. H. Evidence for a direct projection from the postrhinal cortex to the subiculum in the rat. Hippocampus 11, 105–117 (2001).

89. Doan, T. P., Lagartos-Donate, M. J., Nilssen, E. S., Ohara, S. & Witter, M. P. Convergent Projections from Perirhinal and Postrhinal Cortices Suggest a Multisensory Nature of Lateral, but Not Medial, Entorhinal Cortex. Cell Rep. 29, 617-627.e7 (2019).

90. Dalton, M. A., McCormick, C. & Maguire, E. A. Differences in functional connectivity along the anterior-posterior axis of human hippocampal subfields. Neuroimage 192, 38–51 (2019).

91. Vos de Wael, R. et al. BrainSpace: a toolbox for the analysis of macroscale gradients in neuroimaging and connectomics datasets. Commun. Biol. 3, 103 (2020).

92. Chaudhuri, R., Knoblauch, K., Gariel, M.-A., Kennedy, H. & Wang, X.-J. A Large-Scale Circuit Mechanism for Hierarchical Dynamical Processing in the Primate Cortex. Neuron 88, 419–431 (2015).

93. Burt, J. B. et al. Hierarchy of transcriptomic specialization across human cortex captured by myelin map topography. bioRxiv 199703 (2017). doi:10.1101/199703

94. Qin, S. et al. Large-scale intrinsic functional network organization along the long axis of the human medial temporal lobe. Brain Struct. Funct. 221, 3237–3258 (2016).

95. Andrews-Hanna, J. R., Reidler, J. S., Sepulcre, J., Poulin, R. & Buckner, R. L. Functional-Anatomic Fractionation of the Brain’s Default Network. Neuron 65, 550–562 (2010).

96. Libby, L. A., Ekstrom, A. D., Daniel Ragland, J. & Ranganath, C. Differential connectivity of perirhinal and parahippocampal cortices within human hippocampal subregions revealed by high-resolution functional imaging. J. Neurosci. 32, 6550–6560 (2012).

97. Polimeni, J. R., Fischl, B., Greve, D. N. & Wald, L. L. Laminar analysis of 7T BOLD using an imposed spatial activation pattern in human V1. Neuroimage 52, 1334–1346 (2010).

98. Huber, L. et al. High-Resolution CBV-fMRI Allows Mapping of Laminar Activity and Connectivity of Cortical Input and Output in Human M1. Neuron 96, 1253-1263.e7 (2017).

99. Finn, E. S., Huber, L., Jangraw, D. C., Molfese, P. J. & Bandettini, P. A. Layer-dependent activity in human prefrontal cortex during working memory. Nat. Neurosci. 22, 1687–1695 (2019).

100. Braak, H. & Braak, E. Neuropathological stageing of Alzheimer-related changes. Acta Neuropathol. 82, 239–259 (1991).

101. Engel J. J. Mesial temporal lobe epilepsy: What have we learned? Neuroscientist 7, 340–352 (2001).

102. Kim, E. J., Pellman, B. & Kim, J. J. Stress effects on the hippocampus: A critical review. Learning and Memory 22, 411–416 (2015).

103. Lisman, J. et al. Viewpoints: How the hippocampus contributes to memory, navigation and cognition. Nat. Neurosci. 20, 1434–1447 (2017).

104. Bernhardt, B. C. et al. The spectrum of structural and functional imaging abnormalities in temporal lobe epilepsy. Ann. Neurol. 80, 142–153 (2016).

105. Blümcke, I. et al. International consensus classification of hippocampal sclerosis in temporal lobe epilepsy: A Task Force report from the ILAE Commission on Diagnostic Methods. Epilepsia 54, 1315–1329 (2013).

106. Goulas, A., Margulies, D. S., Bezgin, G. & Hilgetag, C. C. The architecture of mammalian cortical connectomes in light of the theory of the dual origin of the cerebral cortex. Cortex 118, 244–261 (2019).

107. Lewis, L., Lepage, C. & Evans, A. An extended MSM surface registration pipeline to bridge atlases across the MNI and the FS/HCP worlds. in (2018).

108. Wagstyl, K. et al. Mapping Cortical Laminar Structure in the 3D BigBrain. Cereb. Cortex 28, 2551–2562 (2018).

109. Wagstyl, K. et al. BigBrain 3D atlas of cortical layers: Cortical and laminar thickness gradients diverge in sensory and motor cortices. PLOS Biol. 18, e3000678 (2020).

110. Waehnert, M. D. et al. Anatomically motivated modeling of cortical laminae. Neuroimage 93, 210–220 (2014).

111. Coifman, R. R. et al. Geometric diffusions as a tool for harmonic analysis and structure definition of data: Multiscale methods. Proc. Natl. Acad. Sci. 102, 7432–7437 (2005).

112. Xiao, Y. et al. An accurate registration of the BigBrain dataset with the MNI PD25 and ICBM152 atlases. Sci. data 6, 210 (2019).

113. Salimi-Khorshidi, G. et al. Automatic denoising of functional MRI data: Combining independent component analysis and hierarchical fusion of classifiers. Neuroimage 90, 449–468 (2014).

114. Griffanti, L. et al. ICA-based artefact removal and accelerated fMRI acquisition for improved resting state network imaging. Neuroimage 95, 232–247 (2014).

115. Pruim, R. H. R. et al. ICA-AROMA: A robust ICA-based strategy for removing motion artifacts from fMRI data. Neuroimage 112, 267–277 (2015).

116. Pruim, R. H. R., Mennes, M., Buitelaar, J. K. & Beckmann, C. F. Evaluation of ICA-AROMA and alternative strategies for motion artifact removal in resting state fMRI. Neuroimage 112, 278–287 (2015).

117. Razi, A., Kahan, J., Rees, G. & Friston, K. J. Construct validation of a DCM for resting state fMRI. Neuroimage 106, 1–14 (2015).

118. Stephan, K. E., Weiskopf, N., Drysdale, P. M., Robinson, P. A. & Friston, K. J. Comparing hemodynamic models with DCM. Neuroimage 38, 387–401 (2007).

119. Friston, K., Mattout, J., Trujillo-Barreto, N., Ashburner, J. & Penny, W. Variational free energy and the Laplace approximation. Neuroimage 34, 220–234 (2007).

120. Friston, K. J. et al. Bayesian model reduction and empirical Bayes for group (DCM) studies. Neuroimage 128, 413–431 (2016).

121. Iglesias, J. E. et al. A computational atlas of the hippocampal formation using ex vivo, ultra-high resolution MRI: Application to adaptive segmentation of in vivo MRI. Neuroimage 115, 117–137 (2015).

122. Desikan, R. S. et al. An automated labeling system for subdividing the human cerebral cortex on MRI scans into gyral based regions of interest. Neuroimage 31, 968–980 (2006).

123. Karapanagiotidis, T. et al. The psychological correlates of distinct neural states occurring during wakeful rest. bioRxiv 1–15 (2019). doi:10.1101/2019.12.21.885772

124. Váša, F. et al. Adolescent tuning of association cortex in human structural brain networks. Cereb. Cortex 28, 281–294 (2018).

125. Alexander-Bloch, A. F. et al. On testing for spatial correspondence between maps of human brain structure and function. Neuroimage 178, 540–551 (2018).

126. Meng, X. L., Rosenthal, R. & Rubin, D. B. Comparing correlated correlation coefficients. Psychol. Bull. 111, 172–175 (1992).

127. Glasser, M. F. et al. The minimal preprocessing pipelines for the Human Connectome Project. Neuroimage 80, 105–124 (2013).

128. Greve, D. N. & Fischl, B. Accurate and robust brain image alignment using boundary-based registration. Neuroimage 48, 63–72 (2009).

129. Murphy, K. & Fox, M. D. Towards a consensus regarding global signal regression for resting state functional connectivity MRI. Neuroimage 154, 169–173 (2017).

